# Roles of Vitamin Metabolizing Genes in Multidrug-Resistant Plasmids of Superbugs

**DOI:** 10.1101/285403

**Authors:** Asit Kumar Chakraborty

## Abstract

Superbug crisis has rocked this world with million deaths due to failure of potent antibiotics. Thousands mdr genes with hundreds of mutant isomers are generated. Small integrons and R-plasmids have combined with F’-plasmids creating a space for >10-20 of mdr genes that inactivate antibiotics in different mechanisms. Mdr genes are created to save bacteria from antibiotics because gut microbiota synthesize >20 vitamins and complex bio-molecules needed for >30000 biochemical reactions of human metabolosome. In other words, mdr gene creation is protected from both sides, intestinal luminal cells and gut bacteria in a tight symbiotic signalling system. We have proposed, to avert the crisis all vitamin metabolizing genes will be acquired in MDR- plasmids if we continue oral antibiotics therapy. Therefore, we have checked the plasmid databases and have detected thiamine, riboflavin, folate, cobalamine and biotin metabolizing enzymes in MDR plasmids. Thus *vit* genes may mobilise recently into MDR-plasmids and are likely essential for gut microbiota protection. Analysis found that *cob* and *thi* genes are abundant and likely very essential than other vit genes.

## Introduction

### MDR bacteria contaminated in air dust and rain water

Life creation is due to chemical evolution and Darwin’s adaptation theory tells the unique way of changes in cellular structure and metabolism in different eco-system. Microscopic unicellular bacteria, fungi, yeast, protozoa, and algae are regulating water and atmosphere circling plant and animal kingdoms (Pathak et al., 1993; Ahmadjian, 2000). A battle among the creatures is maximum at all points where simple bacteria create crisis in the human life creating toxins and other metabolic effects (Grossart et al., 2013; Mckenna, 2013). Microbes are in symbiotic relation with human and 2×10^12^ gut bacteria perform various syntheses of bio-molecules like vitamins, butyrate, unsaturated fatty acids, bile salt and prostaglandin (Le Chatelier et al., 2013). Recently, multidrug-resistant microbes have increased in global water and atmosphere (PM10 particulates increased) posing a threat to human population worldwide (Ibiene et al., 2011; Chakraborty, 2017b). Antibiotics had cured most infections between1940-1990 although gradual increase in drug resistance was detected in many continents as early as 1960 (Reynolds, 1989; Paul et al., 2013). Indeed the drug industry had always run to discover potent derivatives like cefotaxime, ceftrioxane, imipenem and dorripenem in place of old drug like ampicillin and oxacillin (Laxminarayan et al., 2013). Most harmful changes have happened when 2-15 kb R-plasmids and integrons are combined with 62kb F’-plasmids and such MDR conjugative plasmids donate mdr genes into all bacteria more easily by conjugation (Chakraborty, 2016a). Thus, 40% common bacteria in Ganga river water and Bay of Bengal sea water are ampicillin resistant and most bacteria (>95%) have isolated from human and animal are ampicillin and tetracycline resistant being two early mdr genes (*amp* and *tet* that have sequenced in 1965 as plasmid pBR322) and many mdr genes are detected in plasmids of intestinal bacteria (Morten et al., 2009).

### Beta-lactamase genes are strongly diversified

Most notorious MDR gene is β-lactamase gene (accession nos. J01749, X92506, AF227505, AF124204) which hydrolyses lactam ring CO-N bond of penicillins and has diversified into 20 different classes (250-382aa) like blaTEM, blaOXA, blaCTX-M, blaCMY, blaKPC and blaNDM-1etc (Bush & Jacoby, 2010; Chakraborty, 2016b; Stiffler et al., 2015). Class B enzymes are Zn++-dependent β-lactamases that demonstrate a hydrolytic mechanism different from that of the serine β-lactamases of classes A, C, and D. Organisms producing these enzymes usually exhibit resistance to penicillins, cephalosporins, carbapenems, and the clinically available β-lactamase inhibitors. Class C AmpC β-lactamases include CMY-2, P99, ACT-1, and DHA-1, which are usually encoded by *bla* genes located on the bacterial chromosome, although plasmid-borne AmpC enzymes are becoming more prevalent. Moreover, OXA-2, OXA-23, OXA-48 and OXA-51/58 have no sequence similarity but all hydrolyses carbapenems and some inhibitors. The *Klebsiella pneumoniae* New Delhi metallo-β-lactamase-1 (NDM-1) was discovered in 2009 in an Indian patient that could hydrolyse all β-lactums (Mataseje et al., 2016). However, NDM1 outbreaks in England and USA as early as 2010 and were found in plasmid as well as chromosome of *K. pneumoniae, E. coli* and *Acinetobacter baumannii* as well as to lesser extent in *Providentia* sp. and *Enterobacter* sp.(Chakraborty, 2016a).

### Other major mdr genes that inactivate antibiotics

*Str*A/B gene (accession nos. KT225462, LN555650) encodes an enzyme (∼267aa) that acetylates streptomycin and diversified *aac* (accession nos. AB061794, JN596279) and *aph (accession nos.* X01702, U32991) mdr genes acetylate and phosphorylate aminoglycoside antibiotics respectively which then could not able to bind ribosome to kill bacteria (Shaw et al., 1993; Villa et al., 2015). The aminoglycoside adenyl transferase [EC:2.7.7.47] was present in many bacterial plasmids like *Escherichia* (accession nos. HG41719, KJ484637, KM377239), *Klebsiella* (accession nos. KF914286, KF719970), *Salmonella* (accession no. JQ899055), *Acinetobacter* (accession no. KM401411), but also present in some bacterial chromosome as in *Salmonella enterica. Cat* gene (accession no. EF516991) acetylates chloramphenicol and acetylated chloramphenicol could not bind 30S ribosome (Robicsek et al., 2006; Schwarz et al., 2004). *Sul*1/2 genes (accession nos. KM877269, AP012056) have been implicated in sulfamethoxazole resistance (Dallas et al., 1992) and *arr3* gene (accession no. KX029331; protein id. APD70456) ribosylates refamycins which then could not inhibit bacterial RNA polymerase (Chakraborty, 2016a).

### Drug efflux genes are accumulating in large plasmids

Major drug efflux MDR genes include *tet* gene isomers which encode a membrane-bound drug efflux protein (∼400aa) which kicks out tetracycline from bacterial cell cytoplasm and also has been diversified into tetA/B/C (accession nos. X75761, KC590080) (McMurry et al., 1980). Other potential genes are MFS, RND and MATE types drug efflux genes that could kick out drugs in a proton-pump mechanism (Sun et al., 2014). The MexAB-OprM system in *Pseudomonas aeruginosa* has the broadest substrate specificity and contributes to resistance to macrolides, aminoglycosides, sulfonamides, fluoroquinolones, tetracyclines and many β- lactams. The loss of the outer membrane protein (porin) OprD, is associated with imipenem resistance and reduced susceptibility to meropenem. Similarly, tetracycline resistant protein, tetM (accession no. AY466395, protein id. AAS45561) binds tetracycline increasing drug MIC (Croft et al., 2013; Wang et al., 2015; Chakraborty, 2016a).

### Target mutations are prominent mechanism for multi-resistance

The majority of rifampicin-resistant clinical isolates of *M. tuberculosis* harbour mutations in the 507-533 coding region of *rpoB* gene creating an altered β-subunit of the RNA polymerase that could not able to bind refampicin and altered KatG gene (S315T mutation) and −15C/T mutation in the promoter may be important in isoniazid resistance targeting NADH-dependent enoyl-acyl carrier protein (ACP)-reductase that is involved in mycolic acid synthesis. *Van*A gene cluster are involved in the vancomycin resistance in *Enterococcus facium* (Merlo et al., 2015; also see, plasmids pIP501 and pAM_1) and *erm*A/B genes are diverged 23S rRNA methyl transferases that give resistant to macrolides (Harme et al., 2015). This implies that bacteria do change its target genes if new mdr gene synthesis is delayed or very high dose antibiotic is used. So bacteria need multiple gateways to inactivate drug quickly to save gut bacteria (Chakraborty, 2016c).

### Variety of large plasmids in multidrug-resistant bacteria

*Bla*KPC gene was located in many *Klebsiella pneumoniae* large (108-317kb) conjugative plasmids (accession nos. NC_022078, NC_014312 and JX283456) as well as NDM1 gene was also located in conjugative plasmids (accession nos JN420336, AP012055). Other *Escherichia coli* plasmid pNDM_Dok01 and *K. pneumoniae* plasmid pKpANDM-1 have flanked IS*Aba125 elements and T4SS* genes (Hu et al., 2012; Feng et al., 2015; Flach et al., 2015). Folster JP et al (2014) reported a *Vibrio cholera* strain 2012EL-2176 harbouring IncA/C2 plasmid containing *blaCMY-2, blaCTX-M-2, blaTEM-1, floR, aac*(*3)-IIa, str*A/B, *sul1/2, dfrA1, dfrA27, tetA, mphA, mdr-genes* and also resistant to ciprofloxacin due to mutation in *gyrA*(S83I)/*parC*(S85L) as also seen in plasmid pMRV150 (accession no. EU116442) (Carattoli, 2009; Huang et al., 2013; Hermi et al., 2015).

### MDR Genes move into chromosome to increase gene dose

In skin infection, methicillin resistance gene (*mecA*) encoding a penicillin-binding protein and activated with mobile genetic element, the staphylococcal cassette chromosome *mec* (SCC*mec*) has been reported in *Staphylococcus aureus* chromosome which also is associated with *bla, aac, aad* and *sul*1/2 types MDR genes. This implies that to avert the drug crisis, *mdr* genes have been mobilized into chromosome to increase in copy number and in association with multiple *mdr* genes (Chakraborty, 2016b).

### Vitamin metabolizing genes in bacteria

*Escherichia coli* K-12 synthesizes thiamine monophosphate (vitamin B1) de novo from two precursors, 4-methyl-5-(beta-hydroxyethyl) thiazole monophosphate and 4-amino-5- hydroxymethyl-2-methylpyrimidine pyrophosphate. Thiamine monophosphate is then phosphorylated to make thiamine pyrophosphate coenzyme. Five tightly linked genes (thiCEFGH) are involved and the thiC gene product is required for the synthesis of the hydroxymethylpyrimidine and thiE, *thiF, thiG*, and *thiH* gene products are required for synthesis of the thiazole. Rizobium plasmids carry many thi genes with ∼38-69% similarities to *E. coli* enzymes (Vander Horn et al., 1993; Rodionov et al., 2002). RibB gene (protein ids. CDQ55093; ABU50713) of *Klebsiella pneumoniae* has similarity to 3,4-dihydroxy-2- butanone 4-phosphate synthase and linked to ribA (protein id. ABU50715), ribH (protein id. ABU50714), ribE (protein id. ABU50712) of *Photobacterium kishitanii* (accession no. AY849504.2) and ribX is riboflavin transport system permease (protein ids. GAX64964, AUS72405) and ribY riboflavin binding protein (protein id. GBD30789) (Lopez et al., 1987; Dallas et al., 1992; Talarico et al., 1991). The complete aerobic and anaerobic pathways for the de novo biosynthesis of B12 are known including lower ligand or 5,6- dimethylbenzimidazole (Hazra et al., 2015).

We have extensively reviewed the beta-lactamases (Chakraborty, 2016b) and drug acetyl transferases (Chakraborty et al., 2017) in plasmids and integrons and their association in other mdr genes (Chakraborty, 2016a; Chakraborty, 2017c,f). From GenBank data analysis, we have observed an increase in drug efflux genes and IS-elements in MDR conjugative plasmids with reducing the TRA conjugative genes. Which means most of the bacteria have now many types of MDR-plasmids and as we are continuing insult with complex oral antibiotics, other necessary changes are obvious (1/4 genes in MDR-plasmids are unknown function) to avert the serious health crisis of human and bacteria under strong symbiosis (Ame J Drug Deli Ther., 2018, in press). So we have investigated here the nature of vitamin synthesizing genes in large MDR conjugative plasmids to prove that superbug horror and antibiotic horror are synonemous and many changes have occurred in bacterial genomes and plasmids as we have increased multiple complex antibiotics doses since 1960s.

## Materials and Methods

Water from Ganga River was collected at the morning from Babu Ghat (Kolkata-700001) and Howrah Station area. About 100µl of water was spread onto 1.5% Luria Bartoni-agar plate containing different concentration of antibiotics at 2-50µg/ml. MDR bacteria were selected in agar-plate containing ampicillin, streptomycin, chloramphenicol, tetracycline or ciprofloxacin at 50, 50, 34, 20 µg/ml respectively. As imipenem and meropenem resistant bacteria were present low (0.08-0.2 cfu/ml water), a modified method was followed. 2 ml 5x LB media was added into 10 ml River/Sea water at 2-10µg/ml concentration and was incubated 24 hrs to get drug resistant bacteria population (Chakraborty, 2015). Meropenem resistant bacteria further selected on ampicillin, tetracycline, chloramphenicol and streptomycin to get the superbugs. Antibiotics were purchased from HiMedia and stored at 2-50mg/ml at −20°C. Antibiotic papers were also purchased from HiMedia according to CLSI standard. Antibiotic papers are: AT-50 (aztreonam), COT-25µg (Cotrimoxazole), Met-10µg (methicillin), CAZ-30µg (ceftazidime), LOM-15µg (lomofloxacin), VA-10µg (vancomycin), AK-10µg (Amakacin), TGC-15µg (tigecycline), LZ-10µg (linezolid), and IMP-10µg (imipenem).

The plasmid DNA was isolated from overnight culture using Alkaline-Lysis Method (Sambrook et al., 1982; Chakraborty & Hodgson, 1988). 16S rDNA gene colour Sanger’s dideoxy sequencing was performed by SciGenom Limited, Kerala, India (Ausubel et al., 1989). PCR amplification was performed using 1 unit Taq DNA polymerase, 20ng DNA template, 0.25mM dXTPs, 1.5mM MgCl_2_, for 35 cycles at 95°C/30” (denaturation)- 52°C/50”(annealing)-72°C/1.5’ (synthesis). The product was resolved on a 1% agarose gel in 1X TAE buffer at 50V for 2-4 hrs and visualized under UV light and photograph was taken (Chakraborty et al., 1993). The 16S rRNA gene amplification and mdr genes sequencing are performed by conventional methods (Sanger et al., 1977; Chakraborty et al., 1991). NCBI BLAST analysis was performed for bacterial specific gene analysis (www.ncbi.nlm.nih.gov/blast) and data was submitted to GenBank. NCBI databases were retrieved using the BLAST programmes (www.ncbi.nlm.nih.gov/blast) (Johnson et al., 2008).

The complete genes are sequenced in plasmids and were analyzed by Seq-2 programme of BLAST. Multalin protein sequence software was used to get the nature of conserved sequences among metallo-class B β-lactamases (King and Strynadka, 2013). Sometime, diverged sequences are manually cut and paste into align position in MS word so that it is appeared both sequences have similarity (Marchler-Bauer et al., 2017). For retrieving any nucleotide. we type the same at the NCBI port (www.ncbi.nlm.nih.gov/nucleotide or Protein) and to BLAST search to type the accession number for protein or DNA into BLAST port (*https://blast.ncbi.nlm.nih.gov/Blast.cgi?PAGE_TYPE=BlastSearch*)[Bairoch and Apweiler, 2000; Altschul, 1997].

## Result

### Cobalamine biosynthesis genes in MDR-plasmids

Vitamin-B_12_ is a very complex organic molecule and is needed for Haemoglobin protein to carry oxygen and to deliver it to cells and tissues. So we have checked if *cob* genes are present in MDR-plasmids of pathogens. *Shewanella bicestrii* plasmid pSHE-CTX-M (Accession no. CP022359, 193kb) contains a cobalamine biosynthesis protein (protein id. ASK71392) and an acetyl-COA carboxylase biotin carboxy carrier protein subunit (protein id. ASK71473; nt. 109480-110718C) including *mdr* genes like *CTX-M-15, aac3’-IIa, tetA, aph3”-Ib/d*, and *sul1*. Cobalamine biosynthetic protein has also detected in plasmids pKAZ2 and pKHM243 and similar DUF 3150 protein identified in many plasmids of *E. coli, S. enterica, C. fruendii* and uncultured bacterium (protein ids. WP_094198533, WP_065203424, BAS21640 and ALG87157). Plasmid pECAZ155_KPC of *E. coli* (accession no. CP019001; 272kb) has two cobalamine biosynthetic genes [potein ids. AQV87341 (261aa) and AQV87400 (425aa)] in associated with *blaKPC, mphA, sul1, aac6’- II, aph6’-1d, blaCMY-2* and *aac6’-1a mdr* genes and drug efflux genes like *MFS, tetB* and *cmlA* (Chakraborty, 2016).

Plasmid K-109-R (154kb; accession no. KX029331) of *Klebsiella pneumoniae* had 12 *mdr* genes (*bla, sul1/2, strAB, arr3, cat* etc) with no vitamin synthesizing gene but when we analyzed the 25 unknown genes by BLAST, it appeared indeed it had cobalamine synthesizing gene (protein id. APD70537) with 99% similarity to chromosomal sequences of many pathogens (protein ids. EEO11799, AKN19322, ALL42370) as well as plasmid (protein id. ARJ33497) and many DUF3150 domain proteins (protein id. CSO43544). Further, unknown protein (protein id. APD70542) appears a DNA methylase, unknown protein (id. APD70504) may be a DfrA family trimethoprim resistant protein (accession no. KX957972, protein id. APU91748) and unknown protein (protein id. APD70468) is related to NAD-dependent DNA ligase (protein id. SAA90237). Simple interpretation is that many unknown genes are vitamin metabolizing or related and might be assembled in many plasmids with no mdr genes but now quickly moved to MDR-plasmids to fulfil the symbiotic signals from host requiring, vitamins to support human metabolosome and conversely the nutrient for bacterial growth in the gastrointestinal biofilm. Similarly, after BLAST search of hundred of unknown proteins, we found cobT or vWF factor (protein ids. ASM79841 and APZ79706) in *K. pneumonia*e plasmids (accession nos. KY882285, 156kb and KX636095, 335kb) with a linked cobalamine DAF3150 domain biosynthesis protein. vWF factor (protein id. AKN19321) indentified in *Salmonella enterica* MDR plasmid pRH-1238 (accession no. KR091911, nt. 11696-109873) that also contained blaCMY-16, strA/B, floR, sul1/2, aphA6, aac6’-1b, mphA, qacE, blaNDM1 etc deadly *mdr* genes (Villa L, Guerra B et al. 2015). FAD synthase (phospho hydrolase type, nt. 3669-2554) was located in p2964TF plasmid of *K. pneumoniae* (accession no. KT935446) and cobyric acid synthase (cobQ; protein id. ALU64791, nt. 19094-19879) and likely involved in riboflavin and cobalamine biosynthesis respectively (Bueno MF et al. 2016). NemN coproporphyrinogen III oxadase enzyme was acquired by Ensifer (protein id. AHK46764) and Rhizobium (protein id. PJI38304) species sharing 65% similarity.

### Thiamine Biosynthesis genes in MDR-plasmids

Thiamine is vitamin B1 and thiamine pyrophosphate (TPP) is required in many biochemical reactions both for bacteria and human. So we checked the *thi* genes in large conjugative plasmids of pathogens. Large *Enterobacter cloacae* plasmid, p22ES-469 (Accession no. CM008897; 468kb) has thimine biosynthetic genes like *thiH, thiA* and *thiC* (protein ids. PIA01549, PIA01545, PIA01544 respectively) as well as enzymes involved in pantothenate metabolism (protein ids. PIA01523, PIA01562). However, no *bla, sul1, acc* or *aph mdr* genes but drug transporters (protein ids. PIA01505, PIA01751, PIA01752). Plasmid pKOX_R1 contains *thiF* gene (protein id. AFN35065) for thiamine biosynthetic pathway including metal resistant genes and many MFS drug transporters as well as common *mdr* genes like *sul1, aac3’, ANT, aph* and CTX-M-3 β-lactamase. *Rhizobium leguminosarum* plasmid (accession no. CP016287; 125kb) has thiamine biosynthetic genes (*thi*G/S/O/C; Protein ids. ANP89691/2/3/4) and *cobW* gene for cobalamine synthesis (protein id. ANP89796) but no *mdr* gene but drug efflux genes. Large plasmids from same organism (∼1098kb; accession no. CP025013/ CP016287) have similar thiamine biosynthetic genes (protein id. ANP89582; AUW45994, AUW45995) and cobalamine biosynthetic genes (*cob*W; protein id. ANP89796). Such plasmids also have other vitamin biosynthetic genes like nicotinamide amidase (protein id. AUW45731; accession no. CP025013), biotin carboxylase and biotin sulfoxide reductase (776aa; protein id. AUW45720) suggesting absolute necessary signal indeed have generated in bacteria against high dose antibiotics and such plasmids have high numbers of ABC (protein id. AUW46396) and RND/MFS (protein id. AUW46415) transporters but no classical *mdr* genes like *tet, strAB, amp, sul* were found. It appears such plasmid is primitive and *mdr* genes have generated lately and we have detected GNAT family N-acetyl transferase (protein id. AUW46329) which is different from cat, aacC1 or aacA1 families acetylating enzymes. This data has suggested that accumulation of vitamin synthesizing enzymes into plasmids is not accidental and only is increasing in MDR- plasmid now due to complex antibiotic use.

### Folic acid biosynthetic genes in MDR-plasmids

We also found folate and thimine metabolism enzymes in *Sinorhizobium meliloti* mega plasmid, pSymA (accession no. AE006469; 1354kb) like 5,10 methylene tetrahydrofolate reductase (protein id. AAK65825; nt. 1211022-1211975), formoyl tetrahydrofolate deformylase (protein id. AAK65824) and thiamine pyrophosphate binding enzyme (protein id. AAK65851) but no classical *mdr* gene but penicillin binding proteins (protein ids. AAK65894, AAK65893) and many ABC or MFS transporters. This supports our hypothesis that bacteria will acquire more enzymes in the vitamin biosynthetic pathway to avert the action of antibiotics and will preserve the symbiotic relation in the intestine. So how and why such genes are now accumulating in MDR-plasmids is a important observation but needs research to elucidate if our hypothesis that all vitamin synthesizing enzymes will be in MDR- plasmids to avert antibiotic actions on gut microbiota which are protected by symbiotic phenomenon during evolution (Wang et al., 2017).

### Biotin biosynthetic genes in MDR and Non-MDR plasmids

An acetyl-COA carboxylase biotin carboxy carrier protein subunit (protein id. ASK71473; Accession no. CP022359, nt. 109480-110718C) located in *Shewanella bicestrii* plasmids. *Rhizobium leguminosarum* large plasmids (accession no. CP025013/ CP016287) have similar biotin carboxylase and biotin sulfoxide reductase (776aa; protein id. AUW45720) genes and also cobW genes involved in cobalamine biosynthesis. *Ensifer adhaerens* large plasmid pOV14b (1614kb) has 200 diversified ABC transporters and 60 diversified transcriptional regulators and few *mdr* genes (blaOXA-like; protein id. AHK47346), PBP1A (protein id. AHK47149) as well as acrB (protein id. AHK47037) and ermA/QacA (protein id. AHK47565) drug efflux proteins, Search indicated that the plasmid had biotin synthesizing enzymes (bioB, bioD and bioF) at the nt. 983383-981237 (protein ids. AHK47270, AHK4768 and AHK47269 respectively). Thus multiple vitamin synthesizing genes are accumulating in mdr plasmids due to repeated use of complex antibiotics.

### Riboflavin biosynthetic genes in large MDR plasmids

FAD synthase (phospho hydrolase type) was located in p2964TF plasmid of *K. pneumoniae* (accession no. KT935446; nt. 3669-2554). FAD-binding monooxygenase enzyme was reported in *Ensifer* plasmid pOV14b (protein id. AHK47466, nt. 1352625-1351060) as well as a FAD-dependent oxidoreductase (protein id. AHK47620).

### Pyrodoxine biosynthetic gene in MDR-plsmids

A pyridoxine 5’-P oxidase (protein id. AHK47166) was reported in pOV14b large plasmid (accession no. CP007239) of *Ensifer sp.* and related to *Sinorhizobium* enzyme (65% similarity, protein id. WP_058323781).

### Pantothenate biosynthetic gene in large MDR Plasmids

*Enterobacter cloacae* plasmid, p22ES-469 (Accession no. CM008897) has acquired enzymes involved in pantothenate metabolism (protein ids. PIA01523, PIA01562) and also in association with *thi* genes and many *mdr* genes.

### Niacin biosynthetic genes in MDR-plasmids

*Rhizobium leguminosarum* large plasmids have nicotinamide amidase gene (protein id. AUW45731; accession no. CP025013, nt. 271544-272197) involved in niacin metabolism in association of thiG/O/C and cobW genes. NH_3_-dependent NAD^+^ synthetase (nadE1, protein id. AHK46730) was located in Ensifer plasmid with 66% similarity to *Agrobacterium* enzyme (protein ids. CUX02012 and OFE24756) and a acyl coenzyme transferase (protein id. AHK47077) could be related to *Rosebacter denitrificans* (protein id. SFF70718, 77% similarity). NADH-Ubiquinone oxido reductase (protein id. AHK47196) was found in Ensfer sp., large plasmid pOV14b (accession no. CP007239, nt. 854771-858262).

### Many biosynthetic genes in large MDR plasmids

As discussed earlier, we detected many large plasmids have multiple *vit* genes. *Enterobacter cloacae* large plasmid p22ES-469 has many *thi* and *pan* genes (Miranda-Ríos et al., 1997). *Shewanella bicestrii* plasmid pSHE-CTX-M has *cob* and *rib* genes. *Rhizobium leguminosarum* very large plasmid also have multiple *thi* genes as well as *bio* and *cob* genes (accession no. CP025013). This implies that gradual selection of *vit* genes into plasmids have initiated and maintained during antibiotic exposure.

## Discussion

Multi-resistance horrors have intensified worldwide. Indian NAP-AMR (The National Action Plan on Antimicrobial Resistance-2017-2021) has pinpointed five main areas: (i) improving awareness and understanding of AMR through effective education and surveillance, (ii) reducing infection by increasing preventive measures, (iv) reducing the use of antimicrobials in health, food animals and agriculture (v) promoting for AMR research and drug innovations and (v) strengthening India’s leadership on AMR and International collaboration (assessed on October, 2017; Chandy et al., 2014). We thus have studied superbugs at the molecular level. Sadly, study indicated that huge antibiotics use did influenced 2×10^12^ bacteria in the intestine at the molecular level altering vitamin synthesizing capacity and critical *mdr* genes including vitamin synthesizing genes are created in plasmids to save bacteria and human under symbiosis. Indeed high oral dose of ampicillin, streptomycin, ciprofloxacin and tetracycline had killed all gut bacteria between1950-1970 before the introduction of B-complex vitamin capsule and bifidobacterrium probiotic capsule (Feng et al., 2016). Thus it appears multidrug-resistant genes creation is to protect microbiota from repeated doses of antibiotics that we have consumed since 1940s. We detected *blaTEM-1, blaCTX-M*-15, *blaNDM-1* and *aac6’-1b mdr* genes in Ganga River water and Digha sea water and such superbugs are resistant to many drugs like imipenem, meropenem, amikacin and linezolid. We used advanced molecular tools like PCR and DNA sequencing to prove the fact. Further, several high quality research from US Human Microbiome Project (HMP), European Metagenomics of the Human Intestinal Tract (MetaHIT) and others have demonstrated the beneficial functions of the normal gut flora (>35000 species) on health. Symbiotic bacteria contribute to the functional biodiversity in the aquatic world and influence the fitness of the host organisms and microhabitats ecosystem. It has been suggested that LPS, vitamins and butyrate from intestinal bacteria activate luminal cells to secrete interleukins and cytokines that help to synthesis diversified *mdr* genes in conjugative plasmids and chromosome to protect gut microbiota against action of high dose of antibiotics (Gibson et al., 2016). We propose that MDR bacteria will be normal resident of intestine until we stop oral antibiotic use. It is thus G-20 Nations in Germany are united for active research on MDR bacteria to stop superbug horror and WHO has warned to use multiple doses of antibiotics to patients and animals. Many microorganisms are also gathering toxin genes in plasmids and likely very threatening to public. *Bacillus thuringiensis* plasmid pBMB293 (Accession no. CP007615, 294kb) has no *mdr* gene but genes for enterotoxins (protein id. AIM34697), dipterans toxin (protein id. AIM34741) and reverse transcriptase, DNA polymerase β, DNA topoisomerase III and type II secretion system. Similarly, *Bacillus anthrus* plasmid pX01 (accession no. CM002399; 171kb) has toxin gene (protein id. AFL55645, 809aa) and also in pBMB293 plasmid. We need to reduce global toxicity in water and our industries must know the knowledge of contamination of chemicals and heavy metals in water increasing superbugs spread.

## Conclusion

WHO has suggested to follow AMR Action Plan and G-20 leaders and Scientists have agreed to reduce antibiotics use in human, animal, and agricultural land as well as to augment research on novel therapeutics alternate to antibiotics (Villa et al., 2015). We have purified organic phyto-extracts (*Cassia fistula, Suregada multiflora, Syzygium aromaticum, Cinnamomum zeynalicum, etc)* that inhibit Kolkata superbugs and gives a hope for new drug development (Chakraborty, 2015). We slogan, “Come Back to Nature: Save Plants and Use as Medicine”. Many technologies like enzybiotics, phage therapy and gene medicines are under development other than herbal therapy (Kutter et al., 2015). Sadly, household water, sea water, rain water and river water are contaminated greatly with superbugs increasing deadly infections where MRSA *Staphylococcus aureus*, MDR *Acinetobacter baumannii* and NDM1 *Escherichia coli* infections are all antibiotic resistant. Thus we lost our win position using ampicillin, streptomycin, tetracycline, ciprofloxacin, azithromycin, and cefotaxime and imipenem drugs against MDR bacterial infections. The GenBank data analysis has proved our hypothesis that vitamin crisis in human due to antibiotic therapy. We think much analysis of unknown genes in superbug plasmids are necessary and likely NIH (NCBI, USA) should act quickly. Perhaps DNA recombination enzymes like tranposases, resolvases, integrases, topoisomerases and plasmids partition enzymes are good target for development of therapeutics against superbugs. Trully, all famous drug companies are in horror to invest in new antibiotic discovery. But phage therapy (bacteriophages T4/θ/ϕ), enzybiotics (lysin, lysozyme), gene medicines (ribozyme, CASPER-CAS, IL gene therapy) and delivery of toxic drugs efficiently using nano-carriers (fullerenes, DNA-Origami) are frontier medical sciences to regulate the 21^st^ Century drug industry (Dunbar et al., 2018). This work may also through some highlight into combat measure and novel drug design against superbugs and there is no similar report in the Pubmed database (Xu et al., 2014).

**Fig. 1.**
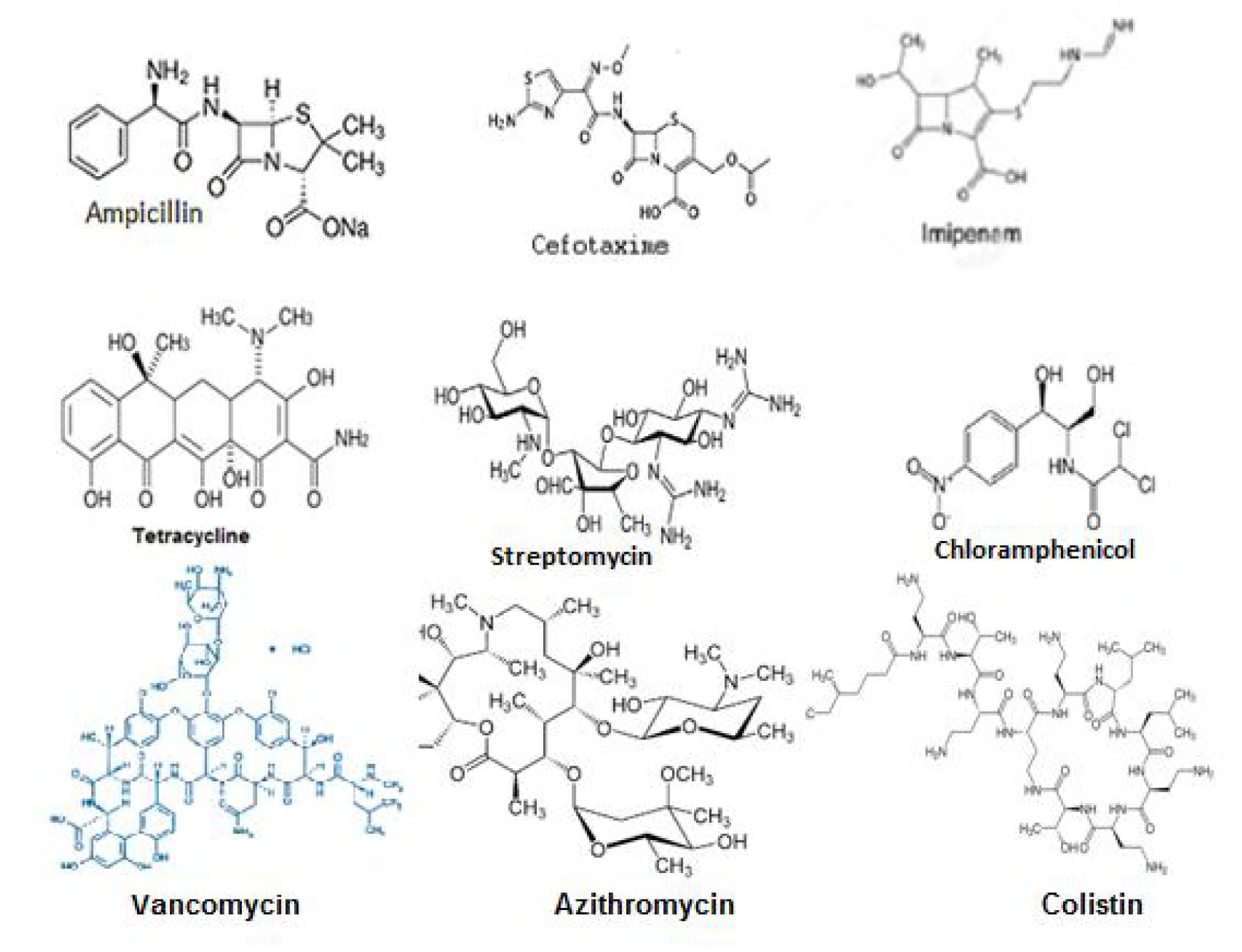
Complex structures of antibiotics that are destroyed by mdr genes and now useless in superbug infections.

**Fig. 2.**
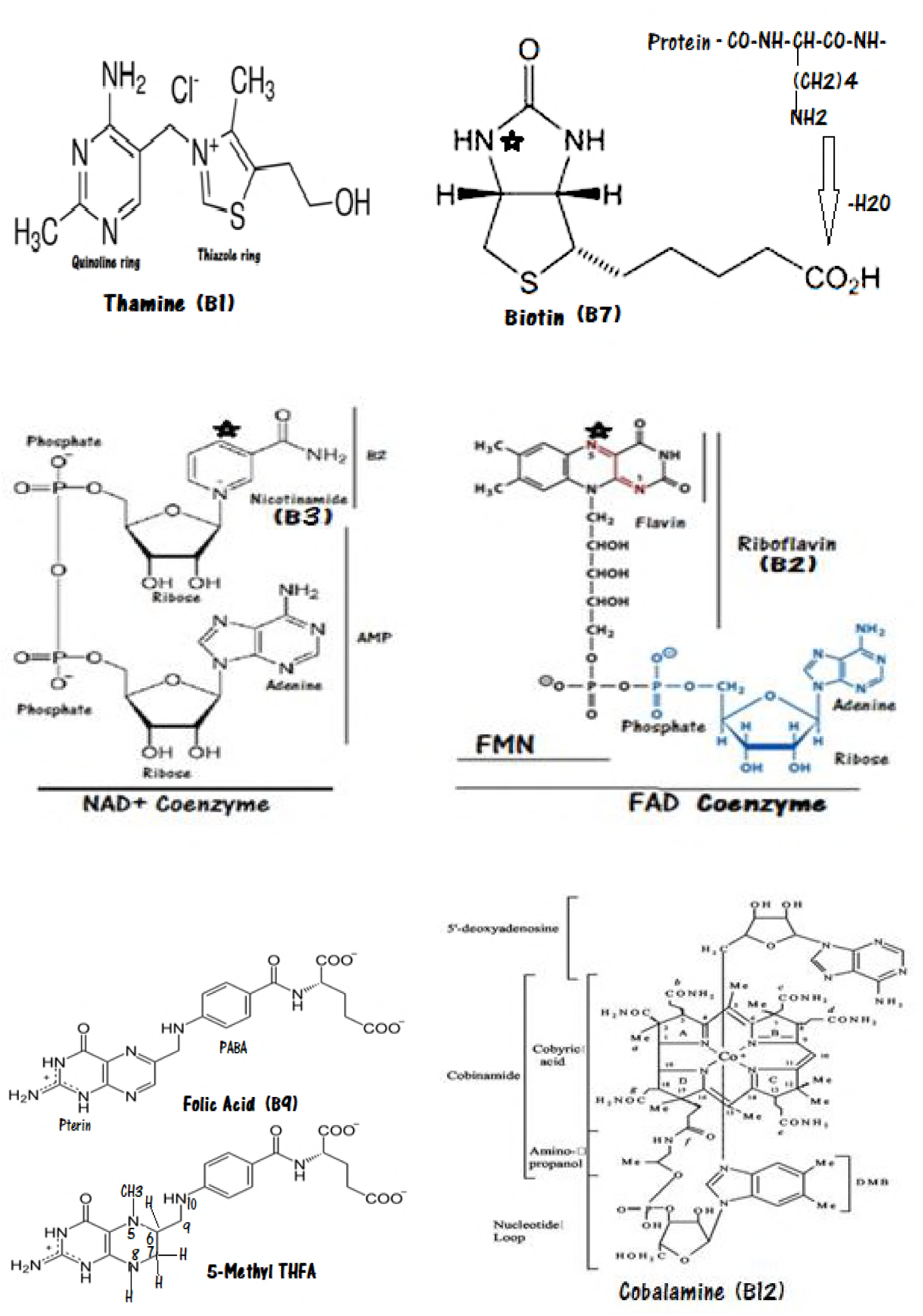
Complex structure of vitamins that intestinal bacteria do synthesis for human.

**Fig. 3.**
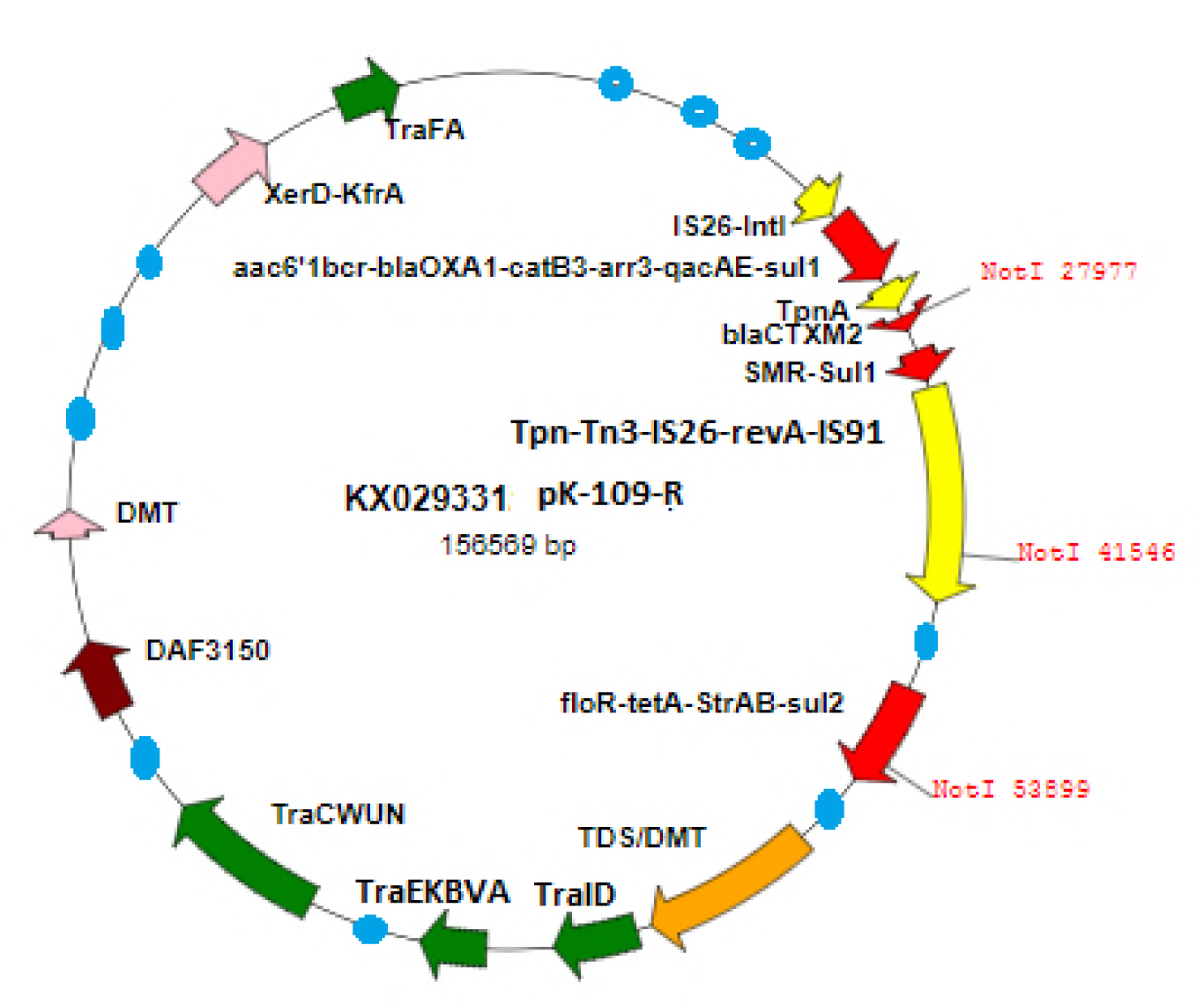
Structure of recent MDR-plasmid pK-109-R (accession no. KX029331). Green is *Tra* genes involved in conjugation; Red is *mdr* genes, Yellow is transposons, Blue is unknown genes and IS elements, and maroon is vitamin synthesizing genes (DAF3150). Restriction enzyme Not1 (octa-cutters) sites are also seen. Such plasmids are capable of donating mdr genes to orther bacteria came to close contact and thus most bacteria became drug resistant.

**Table 1.** Identification of Vitamin Synthesizing Genes in MDR-plasmids with drug efflux and mdr genes. Rizobium plasmids have more drug efflux and ABC transporters but less MDR genes.

| **TABLE-1: Detection of Vitamin metabolizing genes in MDR conjugative plasmids of bacteria** |  |  |  |  |  |
| --- | --- | --- | --- | --- | --- |
| *Vit* genes | PLASMID | Bacteria (Genus) | Accession no | Protein id | Enzyme name and function |
| *thiA* | p22ES-469, 469kb | Enterobacter | CM008897 | PIA01545, 213aa | Thiamine phosphate synthase |
| *thiH* | p22ES-469, 469kb | Enterobacter | CM008897 | PIA01549, 375aa | 2-imino acetate synthase |
| *thiI* | pSymA | Sinorhizobium | AE006469 | AAK65851, 566aa | Thiamine-PP binding protein |
| *thiG* | P1, 125kb<br>pRLN1, 110kb | Rizobium<br>Rizobium | CP016287<br>CP025013 | ANP89691, 257aa<br>AUW45996, 257aa | Catalyses the formation of thiazole. EC: 1.5.3.8. |
| *thiS* | P1, 125kb | Rizobium | CP016287 | ANP89692, 65aa | Ubiquitin-like sulphur carrier protein |
| *thiO* | P1, 125kb | Rizobium | CP016287 | ANP89693, 329aa | FAD-dependent oxidoreductase |
| *thiC* | P1, 125kb<br>pRLN1, 110kb<br>p22ES-469, 469kb | Rizobium<br>Rizobium<br>Enterobacter | CP016287<br>CP025013<br>CM008897 | ANP89694, 607aa<br>AUW45993, 607aa<br>PIA01544, 631aa | Phosphomethyl pyrimidine synthase |
| *thiF* | pKOX_R1, 354kb<br>pC13-2, 104kb | Klebsiella<br>Acinetobacter | CP003684<br>KU549175 | AFN35065, 742aa<br>AMQ95348, 576aa | Thiamine synthetase |
| *thiE* | P2, 181kb<br>pRLN1, 110kb | Vibrio<br>Rizobium | CP022555<br>CP025013 | ASQ38758, 204aa<br>AUW45997, 202aa | Thiamine phosphate pyrophosphorylase |
| *cobW* | P1, 125kb | Rizobium | CP016287 | ANP89796 | Cobalamine biosynthetic protein |
| *cobX* | pKHM-1, 166kb<br>pVPS129, 157kb<br>pRJ119-NDM1, 335kb<br>pECAZ155_KPC, 272kb | Citrobacter<br>Vibrio<br>Klebsiella<br>Escherichia | AP014939<br>KY014464<br>KX636095<br>CP019001 | BAS21640, 425aa<br>ARJ33497, 425aa<br>APZ79705, 425aa<br>AQV87400, 425aa | DUF3150 domain protein. Na_Ca_ex domain synthesizing genes |
| *cobQ* | pECAZ155_KPC, 272kb | Escherichia | CP019001 | AQV87341, 261aa | Cobalamine biosynthetic protein |
| *cobS* | pRJ119-NDM1, 335kb | Klebsiella | KX636095 | APZ79701, 332aa | Cobalamine chelatase |
| *thfR* | pSymA, 135kb | Sinorhizobium | AE006469 | AAK65825, 317aa | 5,10 methylene tetrahydrofolate reductase, EC:1.5.1.20 |
| *thfD* | pSymA, 135kb | Sinorhizobium | AE006469 | AAK65824, 286aa | Formyl tetrahydrofolate deformylase, EC:3.5.1.10 |
| *pdxH* | P2, 181kb | Vibrio | CP022555 | ASQ38911, 211aa | pyridoxamine 5'-phosphate oxidase |
| *bioC* | pCB782, 156kb | Rizobium | CP007068 | AHG48429, 464aa | biotin carboxylase |
| *bioD* | pRLN1, 110kb | Rizobium | CP025013 | AUW45720, 776aa | biotin sulfoxide reductase |
| *bioT* | pSHE-CTX-M, 193kb | Shewanella | CP022359 | ASK71473, 412aa | acetyl-CoA carboxylase biotin carboxy carrier protein |
| *nicA* | pRLN1, 110kb | Rizobium | CP025013 | AUW45731, 217aa | Nicotinamide amidase |

## Acknowledgement

We thank Profs. Amiya Panda and Chandradipa Ghosh of Vidyasagar University; Dr. Sujoy Dasgupta of Bose Institute, Kolkata; Dr. Ananta Ghosh of IIT-Kharagpur and Dr. Ramdhan Mazi of Indian Institute of Chemical Biology, Kolkata for help during the study.

